# Shared and distinct temporal representations of Chinese words across imagined, silent, and overt speech

**DOI:** 10.64898/2026.08.25.747136

**Authors:** Lu Nie, Zitong Lu

## Abstract

How internal speech relates to overt speech remains a fundamental question in speech production: do different forms of speech preserve a common neural representation of the intended word, or does that representation change as speech becomes articulated? We used time-resolved electroencephalography to characterize representations of 10 Chinese words during imagined, silent, and overt speech. Word identity was reliably decodable in all three modes, but its temporal dynamics differed: imagined-speech representations peaked earlier and were less temporally stable, whereas silent and overt speech showed stronger and more sustained representations. Cross-mode decoding revealed word-discriminative information shared across all three mode pairs, with substantially stronger generalization between silent and overt speech. However, direct comparison of word-level representational geometry revealed robust correspondence only between silent and overt speech, indicating that transferable information across modes does not necessarily imply preservation of the broader relational structure among words. Representational similarity analyses further showed distinct visual-form, semantic, and phonetic dynamics across speech modes, with late visual-form and phonetic information contributing uniquely to the geometry shared by silent and overt speech. Controlling for time-matched surface electromyography preserved the overall silent-overt neural correspondence and within-mode phonetic representations, while eliminating the unique phonetic contribution to their shared geometry, suggesting that peripheral articulation accounts for part, but not all, of this common structure. Together, these findings show that imagined, silent, and overt speech share word representations at different levels and suggest that representational geometry and temporal stability are progressively reorganized as internal speech is translated into articulation.

## Introduction

Speech can be generated in markedly different forms. A word may be spoken aloud, articulated without vocalization, or produced entirely in imagination. These modes differ substantially in their physical implementation: overt speech culminates in articulatory movements and their auditory and somatosensory consequences, silent articulation preserves much of the articulatory component while eliminating vocal output, whereas imagined speech can occur without observable articulation. Yet all three modes can instantiate the same intended linguistic content. The relationship among these forms of speech has long been central to theories of inner speech and speech production, which differ in the extent to which internal speech is assumed to preserve the phonological, phonetic, and articulatory detail of overt production (Alderson-Day and Fernyhough, 2015; Oppenheim and Dell, 2010; Perrone-Bertolotti et al., 2014). Behavioral evidence further suggests that inner speech is not simply an impoverished copy of overt speech: the representational detail expressed during internal production can vary depending on whether articulatory movements are permitted (Oppenheim and Dell, 2010). This raises a fundamental question about the organization of speech representations: **to what extent is the neural representation of a word preserved across different modes of speech generation, and to what extent is it transformed as speech becomes increasingly physically instantiated?**

Neuroimaging and electrophysiological studies suggest that different forms of speech are neither fully independent nor neurally equivalent. Functional MRI studies comparing covert and overt production have revealed substantial overlap in the cortical systems recruited by the two modes, together with stronger or additional engagement of regions associated with the motor implementation of speech during overt production (Huang et al., 2002; Palmer et al., 2001; Rosen et al., 2000; Shuster and Lemieux, 2005). Electrophysiological recordings providing complementary evidence at finer temporal scales have further shown overlapping but distinct spatiotemporal activity across speech modes (Pei et al., 2011b; Soroush et al., 2023; Whitford et al., 2017), successful decoding of speech content during both overt and imagined speech (Pei et al., 2011a), and mode-dependent differences in spectral dynamics and representational organization (Proix et al., 2022). Recordings spanning overt, mouthed, and imagined speech likewise reveal systematic changes in speech-related activity as behavioral output is progressively reduced (Soroush et al., 2023), while simultaneous EEG-fMRI demonstrates differences in the temporal and network organization of covert and overt production (Zhang et al., 2024). Together, these findings indicate substantial neural commonality across speech modes alongside mode-dependent differences in their implementation. However, most previous studies have largely characterized neural activity or decoding performance within individual speech modes, with comparatively little direct analysis linking the representational structure across modes. Overlap in regional activation, or the presence of decodable speech information in multiple modes, does not by itself establish that the same words are organized in a common representational format. Moreover, even when related representations are present across modes, they may emerge at different latencies, persist for different durations, or undergo different temporal transformations. Direct cross-mode analyses are therefore needed to determine not only whether word-level representations correspond across imagined, silent, and overt speech, but also how that correspondence unfolds over time.

A further unresolved question concerns what information is shared across speech modes. Word representations are multidimensional, incorporating information related to visual or orthographic form, meaning, and sound. Models of speech production further distinguish interacting processes associated with conceptual and lexical access, phonological encoding, phonetic preparation, and articulation, which unfold dynamically over the course of production (Indefrey, 2011; Indefrey and Levelt, 2004; Liu et al., 2011; Zhang and Damian, 2009). For visually presented words, information about visual form can therefore coexist with semantic and phonological or phonetic information as the system proceeds toward speech production. Studies of Chinese word and character processing have provided evidence that orthographic, phonological, and semantic information can be experimentally dissociated and are associated with overlapping but differentiable neural systems (Cao et al., 2009; Guo and Burgund, 2010; Kuo et al., 2004; Liu et al., 2022; Wu et al., 2012; Zhao et al., 2017). This makes it possible to ask not only whether a word is represented, but which dimensions of that word structure are expressed at different stages of speech generation. Importantly, however, observing the same feature information within two modes does not necessarily mean that this feature contributes to their shared neural structure.

Cross-mode similarity could predominantly reflect semantic information preserved across different forms of production, phonetic information associated with speech planning, visual-form information retained from the presented stimulus, or different mixtures of these dimensions over time. Conversely, a particular feature may be represented reliably within multiple modes but embedded in mode-specific neural patterns that do not generalize across them. A mechanistic account of shared speech representations therefore requires separating two related questions: which features are represented within each speech mode, and which features uniquely contribute to the neural representational structure shared across modes?

An additional challenge is that apparent representational similarity – particularly between silent and overt speech – could partly reflect common peripheral articulatory activity. Surface electromyography (sEMG) provides a sensitive measure of speech-related muscular activity (Stepp, 2012), and articulatory muscle signals can covary systematically with speech production (Schultz and Wand, 2010; Soon et al., 2017). Subtle muscular activity has also been investigated during inner and subvocal speech, although whether such signals consistently carry phonetic information during inner speech remains unresolved (Nalborczyk et al., 2020). This issue is particularly relevant for scalp EEG because craniofacial muscular activity can contribute signals that covary with speech production. Consequently, feature-specific neural structure or similarity between silent and overt speech could in principle be partly attributable to common articulatory activity rather than to shared central representations. Explicitly modeling speech-related sEMG therefore provides a means of testing whether representational effects persist beyond variance associated with peripheral articulation.

Here, we used time-resolved EEG to characterize how representations of Chinese words unfold during imagined, silent, and overt speech. We first asked whether individual words could be reliably distinguished within each speech mode and whether the temporal dynamics and stability of these representations differed across modes. We next tested whether word representations generalized across modes, providing a direct test of shared representational format beyond within-mode decodability. We then used representational similarity analysis (RSA) (Kriegeskorte et al., 2008) to characterize visual-form, semantic, and phonetic information within each mode and to determine which of these feature dimensions uniquely contributed to representational structure shared across modes. Finally, we incorporated sEMG to test whether feature-specific and shared representations, particularly those common to silent and overt speech, could be accounted for by peripheral articulatory activity. Together, these analyses provide a temporally resolved framework for distinguishing which aspects of word representations are shared across different forms of speech generation and which are reorganized as internal representations are translated into articulation.

## Results

### Chinese word representations show distinct temporal dynamics across speech modes

We first asked whether the identity of individual Chinese words could be decoded from EEG activity during imagined, silent, and overt speech, and whether the temporal organization of these representations differed across speech modes. Participants produced each of 10 visually presented single-character Chinese words under the three speech conditions (Figure 1A-B). Time-resolved decoding revealed significant information about word identity in all three modes (Figure 1C). Decoding emerged shortly after stimulus presentation and remained significantly above chance over extended periods in each condition. However, the magnitude and temporal profile of decoding differed markedly across modes. Imagined speech showed a comparatively modest and relatively early increase in decoding accuracy, whereas silent and overt speech exhibited substantially stronger responses that continued to increase later in the trial, with overt speech reaching the highest decoding accuracy.

**Figure 1.**
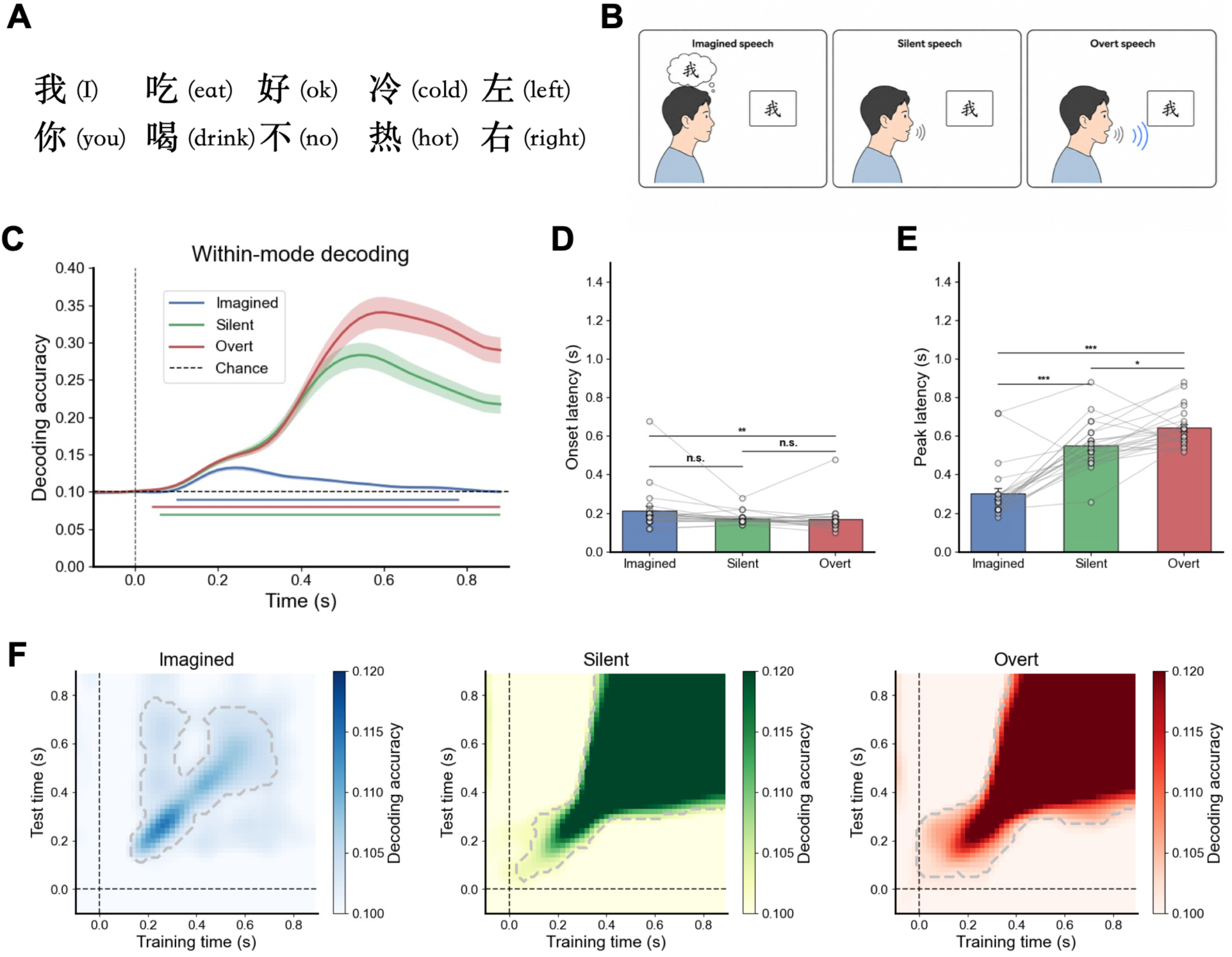
Chinese word representations show distinct temporal dynamics across imagined, silent, and overt speech. (A) The 10 Chinese single-character words used as experimental stimuli, spanning pronouns, verbs, evaluative terms, temperature-related adjectives, and spatial-direction terms; English translations are shown in parentheses. (B) Schematic illustration of the three speech modes. Participants viewed the same visually presented Chinese character and either imagined saying the word without overt articulation (imagined speech), articulated the word silently without vocalization (silent speech), or spoke the word aloud (overt speech). (C) Time-resolved within-mode decoding of word identity for imagined, silent, and overt speech. Curves show the across-participant mean decoding accuracy and shaded regions indicate the SEM. The horizontal dashed line indicates chance-level performance (chance = 0.10), and colored horizontal segments indicate significant above-chance temporal clusters identified using one-sided cluster-based permutation tests (1,000 permutations; cluster-level *p* < 0.05). (D) Onset latency and (E) peak latency of significant word-identity decoding across the three speech modes. Bars show the across-participant mean and SEM; connected points indicate individual participants with valid latency estimates in all three modes. Horizontal annotations indicate Bonferroni-corrected paired comparisons (*p* < 0.05, \**p* < 0.01, \*\**p* < 0.001; *n.s.*, not significant). (F) Within-mode temporal-generalization matrices for imagined, silent, and overt speech. Classifiers were trained at each time point and tested across all other time points; the *x*- and *y*-axes indicate training and test time, respectively. Color indicates mean decoding accuracy, and dashed contours delimit significant above-chance clusters identified using cluster-based permutation tests. Vertical and horizontal dashed lines indicate stimulus onset (0 s).

To quantify these temporal differences, we compared the onset and peak latencies of word-identity decoding across modes (Figure 1D-E). Decoding onset occurred significantly earlier for overt than imagined speech, whereas the onset difference between imagined and silent speech and that between silent and overt speech did not reach significance (Figure 1D). More pronounced differences emerged in peak latency. Imagined speech peaked significantly earlier than both silent and overt speech, and silent speech in turn peaked significantly earlier than overt speech (Figure 1E). Thus, although information distinguishing word identity emerged at broadly comparable early stages, the time at which that information became maximally discriminative shifted progressively later from imagined to silent to overt production.

Within-mode temporal-generalization analyses further revealed different patterns of representational stability across the three modes (Figure 1F). Imagined speech showed a relatively restricted region of above-chance temporal generalization, centered around the period of its early decoding maximum and extending more weakly to later test times. In contrast, silent and overt speech showed broad generalization patterns extending across later portions of the trial. In both articulated conditions, representations emerging from approximately the middle of the trial generalized over an extended range of subsequent test times, with particularly strong and sustained generalization for overt speech. Together, these results show that word identity is represented during all three forms of speech, but that the strength, peak timing, and temporal stability of these representations differ substantially across speech modes.

### Chinese word representations generalize across speech modes, most strongly between silent and overt speech

The presence of decodable word information within each speech mode does not establish whether the underlying discriminative structure is shared across modes. We therefore tested whether a decoder trained to distinguish words in one speech mode could generalize to another. For each pair of modes, we performed temporal-generalization analyses in both transfer directions and averaged the two directional matrices to quantify bidirectional cross-mode generalization (Figure 2).

**Figure 2.**
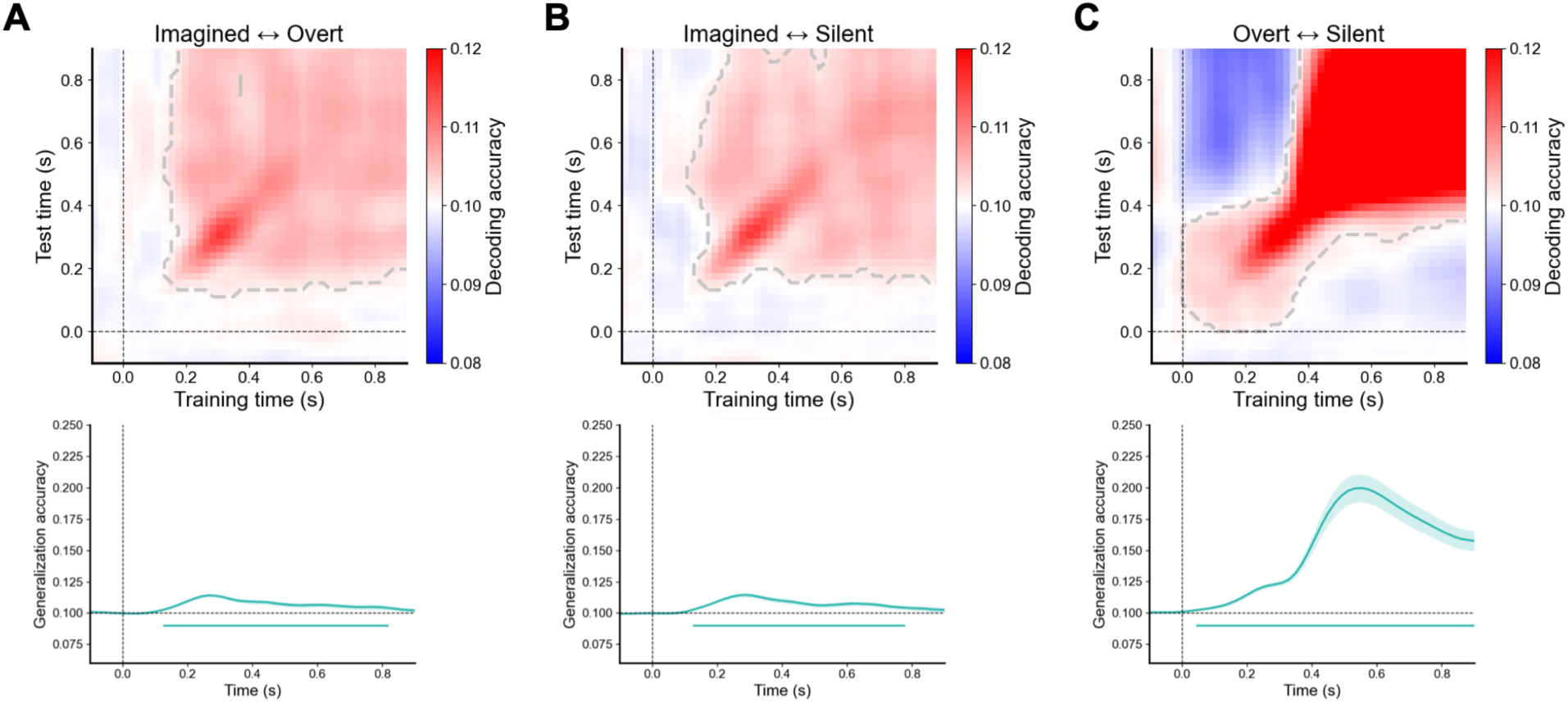
Cross-mode generalization of Chinese word representations is strongest between silent and overt speech. (A-C) Bidirectional cross-mode temporal-generalization matrices for imagined-overt, imagined-silent, and overt-silent speech, respectively. For each participant and mode pair, decoding accuracies from the two opposing transfer directions were averaged element-wise before group averaging. Colors indicate mean cross-mode decoding accuracy. Horizontal and vertical black dashed lines indicate stimulus onset (0 s), and gray dashed contours delimit significant above-chance clusters identified using one-sided two-dimensional cluster-based permutation tests (chance = 0.10; 1,000 permutations; cluster-level *p* < 0.05). (D–F) Corresponding main-diagonal cross-mode decoding time courses for imagined-overt, imagined-silent, and overt-silent speech. Curves show the across-participant mean bidirectional generalization accuracy and shaded regions indicate the SEM. The horizontal dashed line indicates chance-level performance (chance = 0.10), the vertical dashed line indicates stimulus onset, and colored horizontal segments indicate significant above-chance temporal clusters identified using one-sided cluster-based permutation tests (1,000 permutations; cluster-level *p* < 0.05).

Significant cross-mode decoding was observed for all three mode pairs (Fig. 2A-C). Word-discriminative information therefore generalized not only between silent and overt speech, but also between imagined speech and each articulated mode. However, the magnitude of this generalization differed markedly across mode pairs. Imagined-overt and imagined-silent transfer produced relatively modest above-chance decoding, concentrated primarily around corresponding early-to-intermediate time points. By contrast, silent-overt generalization was substantially stronger and extended over a much broader region of training and test times. In particular, representations trained during later silent or overt speech generalized across an extended period in the other mode, indicating sustained correspondence between the discriminative structures expressed during silent and overt production.

The main-diagonal time courses showed the same pattern (Figure 2D-F). Bidirectional generalization between imagined and overt speech and between imagined and silent speech rose modestly above chance before returning toward baseline later in the trial. Silent–overt generalization, in contrast, increased sharply from the intermediate portion of the trial, reached a markedly higher maximum, and remained strongly above chance through the end of the analyzed period. Analyses retaining the two transfer directions separately yielded the same qualitative conclusion: significant generalization was present in both directions for each mode pair, while transfer between silent and overt speech was consistently much stronger than transfer involving imagined speech (Figure S1). These results indicate that some word-discriminative information is shared across all three speech modes, but that silent and overt speech are characterized by a substantially stronger shared discriminative format.

### Feature-specific representational dynamics across speech modes

Having established that word identity was represented within each speech mode and could generalize across modes, we next asked what dimensions of the words were reflected in these neural representations. We constructed three representational models capturing visual-form, semantic, and phonetic relationships among the 10 Chinese words (Figure 3A). The visual-form RDM characterized similarity in the visually presented character forms, the semantic RDM captured relationships in word meaning, and the phonetic RDM characterized similarity in pronunciation-related features.

**Figure 3.**
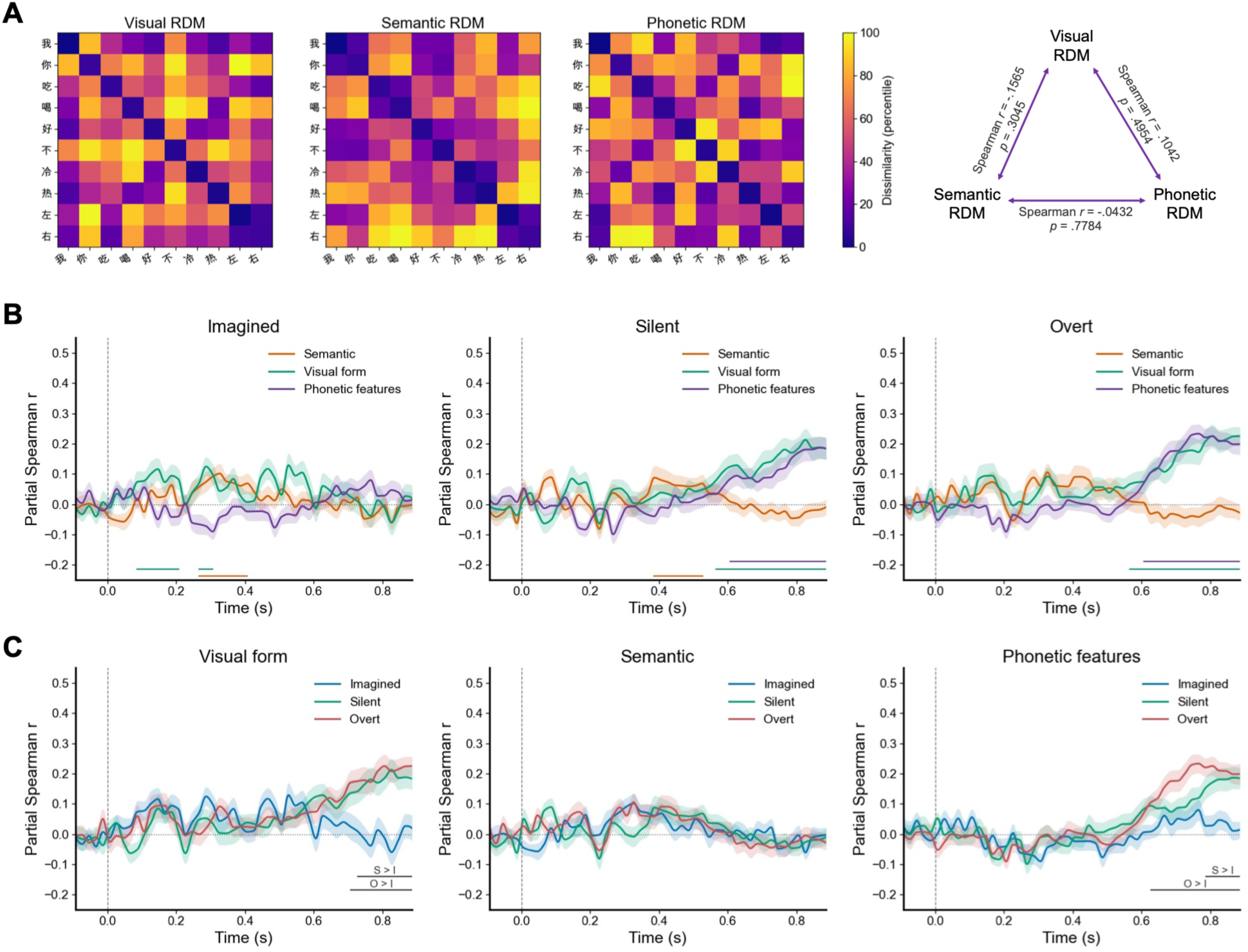
Feature-specific representational dynamics differ across imagined, silent, and overt speech. (A) Three feature-based RDMs corresponding to visual, semantic, and phonetic features. (B) Within-mode partial RSA time courses for imagined, silent, and overt speech. For each speech mode, EEG representational dissimilarity matrices were related to semantic, visual-form, and phonetic-feature models using partial Spearman correlation while controlling for the other two feature models. Curves show the across-participant mean partial correlation and shaded regions indicate the SEM. The horizontal dashed line indicates zero partial correlation, the vertical dashed line indicates stimulus onset (0 s), and colored horizontal segments mark significant positive feature-related temporal clusters identified using one-sided cluster-based permutation tests (1,000 permutations; cluster-level *p* < 0.05). (C) The same partial RSA results regrouped by representational feature to directly compare imagined, silent, and overt speech. Curves and shaded regions again indicate the across-participant mean and SEM. Horizontal bars at the bottom indicate significant pairwise differences between speech modes identified using two-sided cluster-based permutation tests (1,000 permutations; cluster-level *p* < 0.05); labels indicate the direction of the effect (I, imagined; S, silent; O, overt).

We then related the time-resolved EEG representational dissimilarity matrices to the visual-form, semantic, and phonetic models using partial Spearman correlation separately within each speech mode (Figure 3B). The three speech modes showed distinct feature-specific temporal profiles.

During imagined speech, visual-form information was expressed relatively early, followed by a significant semantic contribution at intermediate latencies. In contrast, no sustained late phonetic representation was evident during imagined speech. Silent speech showed a different progression. Semantic information was significantly represented during an intermediate period, whereas visual-form information became prominent later and remained elevated through the end of the epoch. Phonetic information also emerged late in silent speech and increased toward the end of the trial. Overt speech showed a similar late rise in visual-form and phonetic representations, with both features showing sustained significant effects during the later portion of the epoch. In particular, phonetic structure became strongly expressed during late overt production.

Regrouping the same RSA results by representational feature made the cross-mode differences more apparent (Figure 3C). Visual-form representations were broadly comparable across modes earlier in the trial but became significantly stronger in both silent and overt speech than in imagined speech at later time points. Semantic representations showed broadly similar time courses across modes, with no significant pairwise mode differences. Phonetic representations, by contrast, diverged strongly at later latencies: overt speech showed significantly stronger phonetic structure than imagined speech over an extended late interval, and silent speech also exceeded imagined speech later in the epoch. Thus, the transition from imagined to articulated speech was associated particularly with the emergence of strong late visual-form and phonetic representational structure, whereas semantic information showed comparatively less differentiation across speech modes.

We next asked whether the cross-mode generalization observed above reflected preservation of the broader relational geometry among words. To address this question, we directly correlated EEG RDMs across every pair of speech modes and every combination of time points (Figure 4A). In contrast to the cross-decoding results, which showed significant generalization for all three mode pairs, robust cross-mode RDM correspondence was selectively observed between silent and overt speech. Silent-overt representational similarity emerged at intermediate latencies and expanded into a broad cross-temporal region, with particularly strong correspondence between later representations in the two modes. No cluster-corrected cross-temporal RDM similarity was detected between imagined and silent speech or between imagined and overt speech.

**Figure 4.**
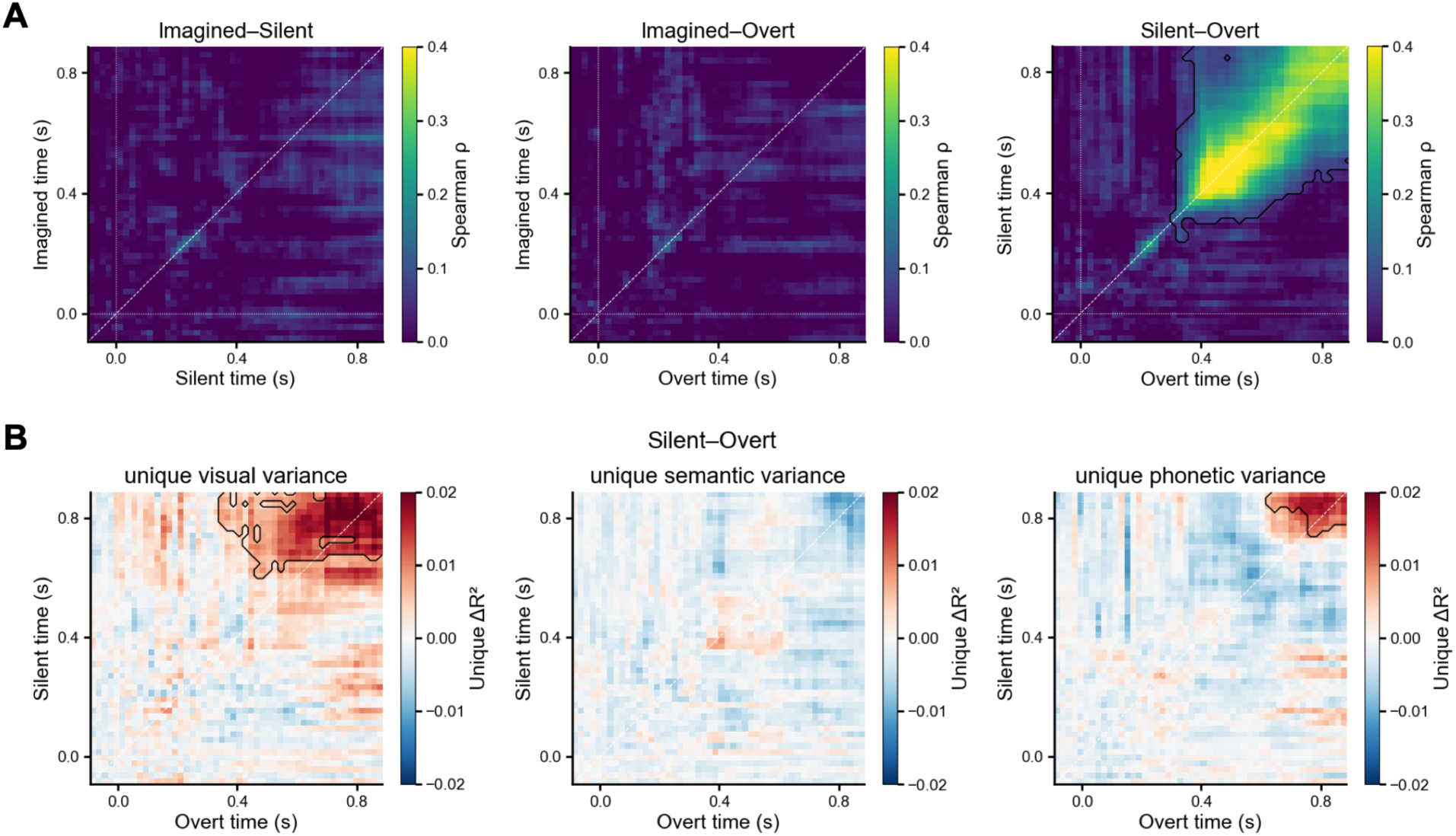
Shared cross-mode representational geometry is selectively expressed between silent and overt speech. (A) Cross-temporal similarity between EEG representational dissimilarity matrices (RDMs) for imagined–silent, imagined–overt, and silent–overt speech. For each pair of speech modes, each matrix element represents the Spearman correlation between the vectorized EEG RDMs at the corresponding pair of time points. Colors indicate the across-participant mean cross-mode RDM similarity. Solid black contours delimit significant positive clusters identified using one-sided two-dimensional cluster-based permutation tests against zero (1,000 permutations; cluster-level *p* < 0.05). Dashed diagonal lines indicate matched time points across the two speech modes. (B) Unique contributions of visual-form, semantic, and phonetic information to the representational geometry shared between silent and overt speech. For each target feature, EEG RDM vectors from both speech modes were first residualized with respect to the other two feature models. Unique ΔR² was then calculated as the difference between the squared residual cross-mode EEG-RDM correlation before and after additionally residualizing both EEG RDMs with respect to the target feature. Positive ΔR² values therefore indicate that removing the target feature reduces cross-mode similarity, consistent with a unique contribution of that feature to the shared geometry. Solid and dashed black contours indicate significant positive clusters, identified using one-sided two-dimensional cluster-based permutation tests (1,000 permutations; cluster-level *p* < 0.05; correction performed separately for each map).

This dissociation indicates that the modest discriminative information shared between imagined speech and the articulated modes was not accompanied by detectable preservation of the broader word-level representational geometry. Silent and overt speech, in contrast, shared both strong discriminative information and a robust relational structure among words. We therefore next asked which feature dimensions uniquely contributed to this silent–overt shared geometry (Fig. 4B). Removing visual-form structure from both EEG RDMs produced a significant reduction in their cross-mode similarity over a broad late region, indicating a unique contribution of visual-form information to the shared silent–overt geometry. Phonetic information also made a significant unique contribution, concentrated at late time points in both modes. In contrast, semantic information did not account for a significant unique component of the shared geometry.

For completeness, we performed the same feature-specific decomposition for the imagined-silent and imagined-overt mode pairs (Figure S2). Neither comparison yielded significant feature-specific clusters. Thus, although imagined speech shared sufficient word-discriminative information with the articulated modes to support cross-mode decoding, we found no evidence that visual-form, semantic, or phonetic structure uniquely accounted for a stable cross-mode representational geometry involving imagined speech. Together, these analyses show that robust word-level representational geometry is selectively shared between silent and overt speech and is supported particularly by late visual-form and phonetic structure.

### Feature-specific and shared representations cannot be explained solely by peripheral articulatory activity

Because silent and overt speech both involve articulatory movements, their feature-specific and shared neural structure could in principle be influenced by common peripheral muscular activity. We therefore repeated the representational analyses while explicitly incorporating time-matched sEMG representational structure.

We first asked whether the feature-specific representations observed within silent and overt speech remained after accounting for peripheral articulation. sEMG itself showed significant word-related representational structure during the later portion of both speech conditions, confirming that peripheral muscle activity carried systematic information about the produced words (Figure 5A). Nevertheless, the principal EEG feature effects remained after sEMG was included as an additional control in the partial RSA. In silent speech, a significant semantic representation was observed at intermediate latencies, whereas visual-form and phonetic representations remained significant later in the trial. Overt speech similarly retained robust late visual-form and phonetic representations after accounting for sEMG. Thus, although peripheral articulatory activity carried word-specific information, it did not fully account for the feature-specific EEG representations observed during silent and overt production.

**Figure 5.**
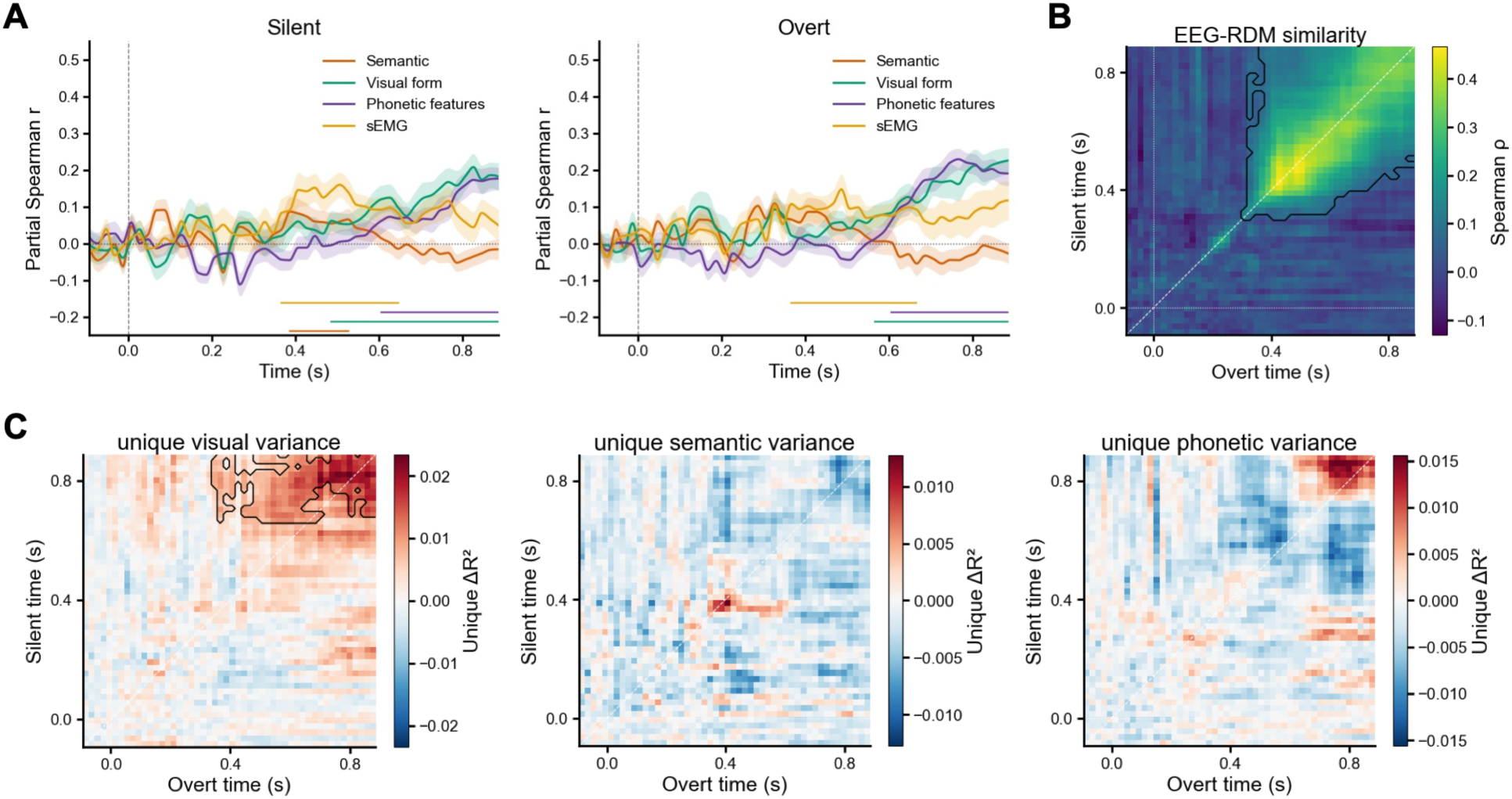
Feature-specific and shared speech representations persist after accounting for peripheral articulatory activity. (A) Within-mode partial RSA for silent and overt speech after including time-matched surface electromyography (sEMG) representational structure in the model. For each speech mode, EEG representational dissimilarity matrices (RDMs) were related to semantic, visual-form, phonetic-feature, and sEMG RDMs using partial Spearman correlation, with the remaining models included as controls for each target model. Curves show the across-participant mean partial correlation and shaded regions indicate the SEM. The horizontal dotted line indicates zero partial correlation, the vertical dashed line indicates stimulus onset (0 s), and colored horizontal segments mark significant positive temporal clusters identified using one-sided cluster-based permutation tests (1,000 permutations; cluster-level *p* < 0.05). (B) Silent–overt cross-temporal partial Spearman similarity between EEG RDMs after simultaneously controlling for the corresponding time-matched silent- and overt-speech sEMG RDMs. Each matrix element reflects the residual cross-mode EEG-RDM similarity at the corresponding pair of time points. The vertical axis indicates silent-speech time and the horizontal axis indicates overt-speech time. Solid black contours delimit significant positive clusters identified using one-sided two-dimensional cluster-based permutation tests against zero (1,000 permutations; cluster-level *p* < 0.05). (C) Unique contributions of visual-form, semantic, and phonetic information to silent–overt shared EEG representational geometry after accounting for the other two feature models and the time-matched silent- and overt-speech sEMG RDMs. For each target feature, EEG RDM vectors from both speech modes were first residualized with respect to the other feature models and both sEMG RDMs. Unique ΔR² was then calculated as the difference between the squared residual cross-mode EEG-RDM correlation before and after additionally residualizing both EEG RDMs with respect to the target feature. Positive ΔR² values indicate that removing the target feature reduces cross-mode similarity, consistent with a unique contribution of that feature to the shared geometry. Solid and dashed black contours indicate significant positive clusters, identified using one-sided two-dimensional cluster-based permutation tests (1,000 permutations; cluster-level *p* < 0.05; correction performed separately for each map).

We next tested whether the broader representational geometry shared between silent and overt speech persisted after simultaneously controlling for the corresponding time-matched sEMG RDMs from both modes. A broad region of significant cross-temporal EEG-RDM similarity remained (Figure 5B), extending across intermediate-to-late time points and closely resembling the silent–overt correspondence observed without sEMG control. The preservation of this cross-mode similarity indicates that the shared word-level EEG geometry between silent and overt speech cannot be explained solely by common peripheral articulatory structure.

Finally, we repeated the feature-specific decomposition of the silent–overt shared geometry while controlling for sEMG from both speech modes (Figure 5C). Visual-form information continued to account for a significant unique component of cross-mode similarity across a broad late temporal region. In contrast, neither semantic nor phonetic information showed a significant unique contribution after sEMG was included in the control set. The loss of the previously observed phonetic unique-variance effect suggests that part of the phonetic structure specifically shared between silent and overt speech overlapped with representational structure present in peripheral articulatory activity. Importantly, however, phonetic information remained significantly represented within the EEG responses of both speech modes, and the overall silent– overt EEG-RDM correspondence remained robust after sEMG control. Together, these results indicate that peripheral articulation contributes to the shared phonetic organization of silent and overt speech but does not fully account for either their feature-specific neural representations or their broader shared representational geometry.

## Discussion

The present study reveals both shared and distinct temporal representations of the same words across imagined, silent, and overt speech. Word identity was decodable in all three modes, yet the representations differed substantially in their strength, timing, and temporal stability. Word-discriminative information also generalized across every pair of speech modes, with markedly stronger transfer between silent and overt speech. Importantly, however, this cross-mode sharing depended on the level of representation considered: robust correspondence of the broader word-level representational geometry was detected only between silent and overt speech. Feature-based analyses further showed prominent late visual-form and phonetic representations in these two articulated modes, both of which contributed to their shared geometry before peripheral articulation was modeled. After controlling for time-matched sEMG, the overall silent–overt geometry and within-mode phonetic representations remained, whereas the unique phonetic contribution to the shared geometry did not. Together, these findings suggest that imagined, silent, and overt speech are neither independent states nor scaled versions of a single neural representation. Instead, they share partially overlapping word information whose temporal organization and representational structure change as speech becomes increasingly articulated.

The distinct temporal profiles across modes extend previous evidence that covert and overt speech differ in more than response magnitude. fMRI studies have demonstrated substantial overlap in the cortical systems recruited by covert and overt production, alongside stronger engagement of motor-related regions during overt speech (Palmer et al., 2001; Shuster and Lemieux, 2005). Electrophysiological studies further show mode-dependent differences in neural dynamics: ECoG recordings reveal overlapping but distinct spatiotemporal activity during covert and overt word production (Pei et al., 2011b; Zhang et al., 2024), while scalp EEG and simultaneous EEG-fMRI identify differences in the timing and organization of speech-related responses across modes (Bowers et al., 2018; Zhang et al., 2024). Our results extend these observations to the dynamics of multivariate word representations. Imagined speech reached its decoding maximum earlier and showed relatively restricted temporal generalization, whereas silent and especially overt speech developed stronger representations that remained stable across broader temporal intervals. This pattern is consistent with speech-production frameworks in which lexical, phonological, phonetic, motor, and sensory processes interact dynamically as production unfolds (Indefrey, 2011; Indefrey and Levelt, 2004). Rather than being a weaker version of overt production, imagined speech therefore appears to follow a distinct representational trajectory, while increasing articulatory implementation is associated with a more sustained neural state. The present EEG data, however, do not allow individual time periods to be assigned uniquely to specific production stages.

A central implication of our findings is that “shared representation” should not be treated as a single property. Previous fMRI and intracranial studies have established substantial overlap in the neural substrates supporting different forms of speech. In particular, Soroush et al., 2023 reported a nested organization across overt, mouthed, and imagined speech, with neural sites relevant to lower-output modes forming subsets of those recruited during more behaviorally expressed production. Our results reveal an additional distinction at the representational level. Cross-mode decoding demonstrated that some word-discriminative information generalized across all three mode pairs, including imagined speech. Yet robust correspondence of the full word-level RDM emerged only between silent and overt speech. Cross-decoding requires only that some discriminative dimensions be preserved across conditions, whereas RDM correspondence places a stronger requirement on preservation of the relational structure among the represented items. Thus, imagined speech may retain a shared subspace sufficient for word-level transfer while reorganizing other dimensions of the representation. Silent and overt speech, by contrast, share a substantially richer representational format. This distinction complements intracranial evidence that imagined and overt speech differ in both spectral organization and the expression of articulatory and phonetic information (Proix et al., 2022). More broadly, neural overlap, transferable information, and preserved representational geometry should be regarded as related but separable levels of cross-mode commonality.

The feature analyses further clarify what changes as speech becomes articulated. Semantic information was represented in all three modes and showed relatively limited differentiation across them, whereas the strongest mode differences occurred in late visual-form and phonetic representations. Silent and overt speech developed pronounced late phonetic structure that was weak or absent during imagined speech, consistent with evidence that articulatory implementation increases the phonetic specificity of internal speech (Oppenheim and Dell, 2010) and with intracranial findings of distinct phonetic and articulatory organization across imagined and overt production (Pei et al., 2011a; Proix et al., 2022). The late visual-form effects were less expected but may be especially relevant for visually presented Chinese characters, for which visual/orthographic, phonological, and semantic representations interact closely (Liu et al., 2022; Wu et al., 2012). Orthographic information can also remain accessible during Chinese spoken-word production (Zhang and Damian, 2009). The visual-form contribution to silent–overt shared geometry therefore raises the possibility that stimulus-form information remains coupled to later production-related representations. This interpretation should remain cautious, however: our visual model captures similarity in image-derived character form and cannot be equated directly with an abstract orthographic code or localized to visual cortex. Future experiments using auditory cues or independently manipulating visual similarity and linguistic identity will be important for determining the source of this late effect.

The sEMG analyses provide an important constraint on the interpretation of the silent–overt similarity. Peripheral muscle activity itself contained reliable word-related representational structure, confirming that common articulation could contribute systematically to apparent cross-mode correspondence. Nevertheless, controlling for time-matched sEMG did not eliminate late phonetic representations within silent and overt EEG, nor did it abolish their broad cross-temporal representational similarity. Peripheral articulation therefore cannot fully explain the shared EEG organization of the two modes. At the same time, the unique phonetic contribution to the silent–overt shared geometry was no longer significant after sEMG control, whereas the visual-form contribution remained. This pattern suggests that peripheral articulatory activity contributes specifically to part of the phonetic structure shared between silent and overt speech, while the broader neural correspondence extends beyond measured muscle activity. This distinction is more informative than treating sEMG simply as an artifact control: it indicates that peripheral and central speech-related representations overlap, but are not equivalent.

Taken together, the results favor a graded but nonuniform relationship among imagined, silent, and overt speech. Imagined speech preserves a component of word identity that generalizes to articulated production, yet its representations are temporally more restricted and do not exhibit the robust global geometry observed between silent and overt speech. Silent speech, despite lacking vocal output, more closely resembles overt speech in temporal stability, feature composition, and relational structure. Articulation may therefore mark an important transition in the organization of internal speech representations. This account is compatible with flexible theories of inner speech, in which representational detail varies with the extent of motor implementation (Alderson-Day and Fernyhough, 2015; Oppenheim and Dell, 2010), but it also suggests that the transition from imagined to overt production cannot be reduced to a simple increase in representational strength. Different aspects of the neural code—discriminative information, overall geometry, and feature-specific structure—change to different degrees across modes.

Several limitations should be considered. First, the three speech modes were presented in a fixed rather than counterbalanced order, so practice, stimulus familiarization, fatigue, or other sequence effects cannot be completely separated from mode. A simple facilitation account, however, does not readily explain the observed pattern. The stimuli were highly familiar Chinese characters, each item was already repeated extensively within each condition, and the strongest between-mode changes were feature- and time-specific rather than a general enhancement in later conditions. In particular, semantic representations did not show the systematic increase that might be expected from broad stimulus facilitation, whereas late visual-form and phonetic effects were selectively enhanced. Nevertheless, future studies should counterbalance mode order.

Second, the representational space contained only 10 words. The large number of repetitions supported stable time-resolved estimates, but larger and more systematically constructed vocabularies will be necessary to test the generality of the observed geometry and more completely dissociate feature spaces. Third, the use of visually presented single Chinese characters limits generalization to other languages, connected speech, and speech generated without an external visual cue. Finally, scalp EEG provides high temporal resolution but limited anatomical specificity; combining these analyses with MEG, fMRI, or intracranial recordings could reveal where shared and mode-specific representational geometries arise.

In summary, speech representations are shared across imagined, silent, and overt production, but at different levels and to different degrees. Word-discriminative information transfers across all three modes, whereas a robust and temporally sustained word-level geometry is most evident between silent and overt speech. The transition toward articulation is accompanied by stronger late visual-form and phonetic organization, with peripheral articulation accounting for part, but not all, of the shared phonetic structure. These findings suggest that internal and overt speech are linked through partially shared representations that are progressively reorganized as linguistic intentions are translated into articulated speech.

## Methods

### Dataset and experimental paradigm

The EEG and surface electromyography (sEMG) data were obtained from an openly available multimodal speech dataset deposited in the Science Data Bank (SCIDB repository: https://cstr.cn/31253.11.sciencedb.24416, Zhao et al., 2025). The present analyses included 30 participants who completed imagined, silent, and overt speech tasks using 10 high-frequency Chinese characters: 我 (I/me), 你 (you), 吃 (eat), 喝 (drink), 好 (good), 不 (no/not), 冷 (cold), 热 (hot), 左 (left), and 右 (right). In the overt condition, participants read the presented character aloud; in the silent condition, they performed the corresponding articulatory movements without producing sound; and in the imagined condition, they read the character internally without overt vocalization. Each character was repeated 50 times within modality-specific blocks, which were completed in the fixed order imagined → silent → overt. Full details of the participants, stimuli, experimental setup, and procedure are provided in Zhao et al., 2025.

### EEG and sEMG acquisition and preprocessing

EEG was recorded with a 64-channel NeuroScan SynAmps2 system at 1,000 Hz, and six-channel facial sEMG was recorded concurrently at 1,000 Hz from speech-related muscle groups. EEG and sEMG systems received synchronized stimulus-trigger codes, enabling time-matched multimodal analyses. Following the preprocessing pipeline described in Zhao et al., 2025, EEG data were cleaned in EEGLAB using bad-channel interpolation, trial rejection, common-average re-referencing, 0-120 Hz band-pass filtering, 50-Hz notch filtering, down-sampling to 256 Hz, baseline correction, and ICA-based artifact removal. sEMG data were band-pass filtered from 1 to 300 Hz, notch filtered at 50 Hz, and segmented using the corresponding EEG event labels. Both modalities were epoched from -0.1 to 0.9 s relative to stimulus onset. For the present decoding and representational analyses, the preprocessed EEG epochs were further resampled to 250 Hz before construction of the time-resolved analysis windows. Further details of data acquisition, synchronization, and preprocessing are available in Zhao et al., 2025.

### EEG word-identity decoding

Within-mode word-identity decoding was performed separately for imagined, silent, and overt speech in 30 participants. The classifier discriminated among 10 Chinese word conditions, corresponding to a chance accuracy of 0.10. Epoched EEG data were cropped from -0.1 to 0.9 s and resampled to 250 Hz. EEG data were arranged as trials × channels × time, and trials were averaged into pseudotrials of four trials before classification. A linear support vector machine was trained separately within non-overlapping five-sample windows (20 ms), with an identical five-sample step. Decoding used stratified five-fold cross-validation with 10 repetitions. Subject-level decoding time courses were obtained before group-level aggregation.

The decoding pipeline was implemented by NeuroRA (Lu and Ku, 2020). Group decoding curves were calculated as the arithmetic mean of the smoothed subject-level accuracies, with shaded regions indicating the between-participant standard error of the mean (SEM). Statistical inference was performed on the same subject-level curves using a one-sample sign-flip cluster-permutation test against chance accuracy (chance = 0.1). Samples exceeding a one-sided cluster-forming threshold of *p* < .05 were grouped into temporally contiguous clusters, and cluster mass was defined as the sum of t values within each cluster. Statistical significance was assessed against the permutation distribution of the maximum cluster mass using 1,000 permutations, a cluster-level threshold of *p* < .05, a positive tail.

### Within-mode temporal generalization

Temporal-generalization decoding was performed separately within each speech mode by training the classifier at each 20-ms EEG window and testing it at every other time window, yielding a train-time × test-time decoding matrix for each participant. The analysis used the same 10-class classification problem, four-trial pseudotrial averaging, stratified five-fold cross-validation, 10 repetitions, training-fold standardization, and absence of PCA as the within-mode time-resolved decoding analysis. Group temporal-generalization maps were obtained by averaging subject-level accuracy matrices and were retained on the original decoding-accuracy scale, with chance accuracy equal to 0.10.

### Cross-mode decoding

Cross-mode decoding tested all six directional transfers among imagined, silent, and overt speech. For each participant and pair of speech modes, trial counts were balanced across the two modes using a participant-specific random seed derived from the participant identifier. A classifier was trained on one speech mode and tested on another across all combinations of training and testing times. The saved analysis used 10 word classes, four-trial pseudotrial averaging, five-sample (20-ms) windows with five-sample steps, 10 repetitions, feature standardization, and principal-component analysis retaining 95% of the variance in the training data. The six directional transfer analyses comprised imagined-to-silent, silent-to-imagined, imagined-to-overt, overt-to-imagined, silent-to-overt, and overt-to-silent decoding. Directional results are reported in Figure S1. For the main bidirectional analysis in Figure 2, the two opposite directional maps for each pair of speech modes were combined at the participant level by elementwise arithmetic averaging.

Thus, for modes A and B,

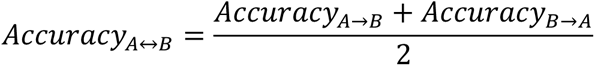

This bidirectional measure summarizes the existing directional transfer results and does not constitute a separate classifier analysis.

### EEG representational dissimilarity matrices

Neural representational dissimilarity matrices (RDMs) were constructed separately for each participant and speech mode. The 10 word conditions were 我, 你, 吃, 喝, 好, 不, 冷, 热,左, and 右. EEG epochs were cropped from -0.1 to 0.9 s and resampled to 250 Hz. For each condition, all available trials were averaged before RDM construction. The resulting condition-specific EEG patterns contained 60 channels.

For each five-sample (20-ms) temporal window, neural dissimilarity was quantified using correlation distance between the channel-by-time activity patterns for every pair of word conditions. Windows advanced in five-sample steps, yielding 50 neural RDMs of size 10 × 10 for each participant and speech mode.

### Feature model RDMs

Three fixed 10 × 10 model RDMs were constructed to capture visual-form, semantic, and phonetic-feature similarity among the 10 words.

The visual-form RDM was derived from activations in the second convolutional layer of an ImageNet-pretrained AlexNet (Krizhevsky et al., 2017). Experimental character images underwent the preprocessing required by the pretrained network, and activation patterns were flattened for each stimulus. Pairwise cosine distance between stimulus activation vectors was used to construct the 10 × 10 visual-form RDM.

The semantic RDM was derived from 200-dimensional Tencent AI Lab Chinese word and phrase embeddings. Embeddings were available for all 10 experimental words, and pairwise cosine distances between embedding vectors were used to construct the semantic RDM.

The distributed phonetic-feature RDM represented the pronunciation structure of each Mandarin monosyllable using a refined 92-dimensional distributed feature vector. Each syllable was decomposed into four segmental slots – onset, glide, nucleus, and coda – together with lexical tone. Each of the four segmental slots was represented using the same 22-dimensional phonetic-feature encoding, yielding 88 segmental dimensions, and lexical tone was represented using a four-dimensional one-hot vector. Concatenating these components yielded a 92-dimensional representation for each word. The feature scheme was based on previous approaches to representing phonological and pronunciation structure in Chinese word processing (Yang et al., 2009; Zhao et al., 2017). Pairwise cosine distances between the 92-dimensional vectors were then used to construct the 10 × 10 distributed phonetic-feature RDM. This model is referred to as Phonetic features in the publication figures and Results.

### Within-mode partial RSA

Within each participant, speech mode, and temporal window, the upper-triangular dissimilarities from the EEG and model RDMs were extracted and rank-transformed. Partial Spearman correlation was then used to quantify the association between the EEG RDM and each target feature RDM while controlling for the other two feature RDMs (Lu and Golomb, 2025). Thus, visual-form information was assessed while controlling semantic and phonetic-feature similarity, semantic information while controlling visual-form and phonetic-feature similarity, and phonetic-feature information while controlling visual-form and semantic similarity.

The resulting without-sEMG-control analysis produced an array organized as 3 speech modes × 30 participants × 3 features × 50 time points. Differences between speech modes were assessed using within-participant difference curves for each feature.

### Cross-mode EEG-RDM similarity

To quantify shared representational geometry across speech modes, cross-temporal EEG-RDM similarity was calculated separately for the imagined–silent, imagined–overt, and silent–overt mode pairs. For each participant, a cell at time coordinates (*t_A_*, *t_B_*) represented the Spearman correlation between the 45 upper-triangular dissimilarities of the EEG RDM from mode A at time *t_A_* and those from mode B at time *t_B_*. This yielded a 50 × 50 cross-temporal similarity matrix for each participant and mode pair.

### Feature-specific unique contribution to shared geometry

To determine which feature dimensions contributed uniquely to representational geometry shared across speech modes, we used the residualization-based procedure implemented in the Figure 4 analysis. For each participant and cross-mode time pair, the ranked EEG dissimilarity vector from each mode was separately regressed onto an intercept and a specified set of feature-model RDM vectors.

For a target feature, *r_without_* was defined as the correlation between the two residual EEG vectors after controlling for the two non-target feature RDMs. The target feature was then added to the residualization model for both speech modes, and *r_with_* was calculated from the resulting pair of residual EEG vectors. The unique contribution of the target feature was quantified as *r^2^_without_* - *r^2^_with_*. This measure therefore quantifies the reduction in squared residual cross-mode EEG-RDM similarity associated with additionally accounting for the target feature. Positive values indicate that accounting for the target feature reduces the squared similarity between the residual EEG representational geometries, whereas negative values are mathematically possible.

Separate unique-contribution maps were calculated for visual form, semantics, and phonetic features. In each analysis, the other two feature models served as controls.

### sEMG representational control analyses

Surface electromyography (sEMG) RDMs were constructed separately for silent and overt speech using the same 10 word conditions. Six sEMG channels were included. As for the EEG RDMs, pairwise representational dissimilarity was quantified using correlation distance. sEMG data were evaluated in five-sample windows with five-sample steps, yielding 50 temporally resolved RDMs. Each sEMG time bin was aligned to the corresponding EEG-RDM time-bin index to provide a time-matched measure of peripheral articulatory representational structure.

### Within-mode RSA with sEMG control

The within-mode sEMG-control analysis focused on silent and overt speech, the two conditions in which participants performed articulatory/vocalization movements. For each participant and time window, four candidate RDMs were considered: semantic, visual form, phonetic features, and the participant- and time-matched sEMG RDM. Partial Spearman correlation was calculated for each target RDM while controlling the remaining three predictors. Consequently, each neural feature estimate quantified visual-form, semantic, or phonetic-feature similarity in the EEG after accounting for the other two linguistic feature models and the concurrent sEMG representational structure.

### Silent–overt shared geometry controlling sEMG

To assess cross-mode shared EEG geometry beyond time-matched peripheral articulatory structure, silent–overt cross-temporal EEG-RDM similarity was recalculated while controlling the corresponding sEMG RDMs. At each silent-overt time pair, the silent EEG RDM was controlled for the silent sEMG RDM at the corresponding silent-speech time point, and the overt EEG RDM was controlled for the overt sEMG RDM at the corresponding overt-speech time point. The correlation between the resulting residual EEG geometries provided the sEMG-controlled cross-mode similarity measure. The three visual-form, semantic, and phonetic-feature RDMs were not additionally controlled in this analysis.

### Feature-specific unique contributions after sEMG control

For the Figure 5 unique-contribution analyses, the residualization framework used for Figure 4 was extended by incorporating the time-matched sEMG RDMs. For each target feature, (*r*_without_) was calculated after separately residualizing the silent and overt EEG-RDM vectors against the two non-target feature models together with the time-matched silent and overt sEMG representational structure. The target feature was then added to the residualization model to obtain (*r*_with_). Unique contribution remained defined as *r*^2^_without_ − *r*^2^_with_.

These analyses therefore quantify feature-specific contributions to silent–overt shared EEG geometry after accounting for the measured, time-matched sEMG representational structure. They do not imply that peripheral articulatory activity makes no contribution to the EEG signal or to speech representations.

### Statistical inference

All group-level inferential analyses treated participant as the sampling unit. One-dimensional cluster tests controlled family-wise error across time within each tested curve, whereas two-dimensional cluster tests controlled family-wise error across connected cells within each train-time × test-time or cross-mode time × time matrix. Other specific statistical details are shown in the corresponding figure caption.

## Data Availability

The EEG and sEMG data used are available as open data via the Science Data Bank (SCIDB) repository: https://cstr.cn/31253.11.sciencedb.24416 (Zhao et al., 2025). In addition, we obtained corrected versions of a subset of the experimental data directly from the dataset authors and used these corrected data in the present analyses. Code for all the analyses in this paper are available at https://github.com/ZitongLu1996/CNWordsSpeechRep_EEG.

## Supplementary

**Figure S1.**
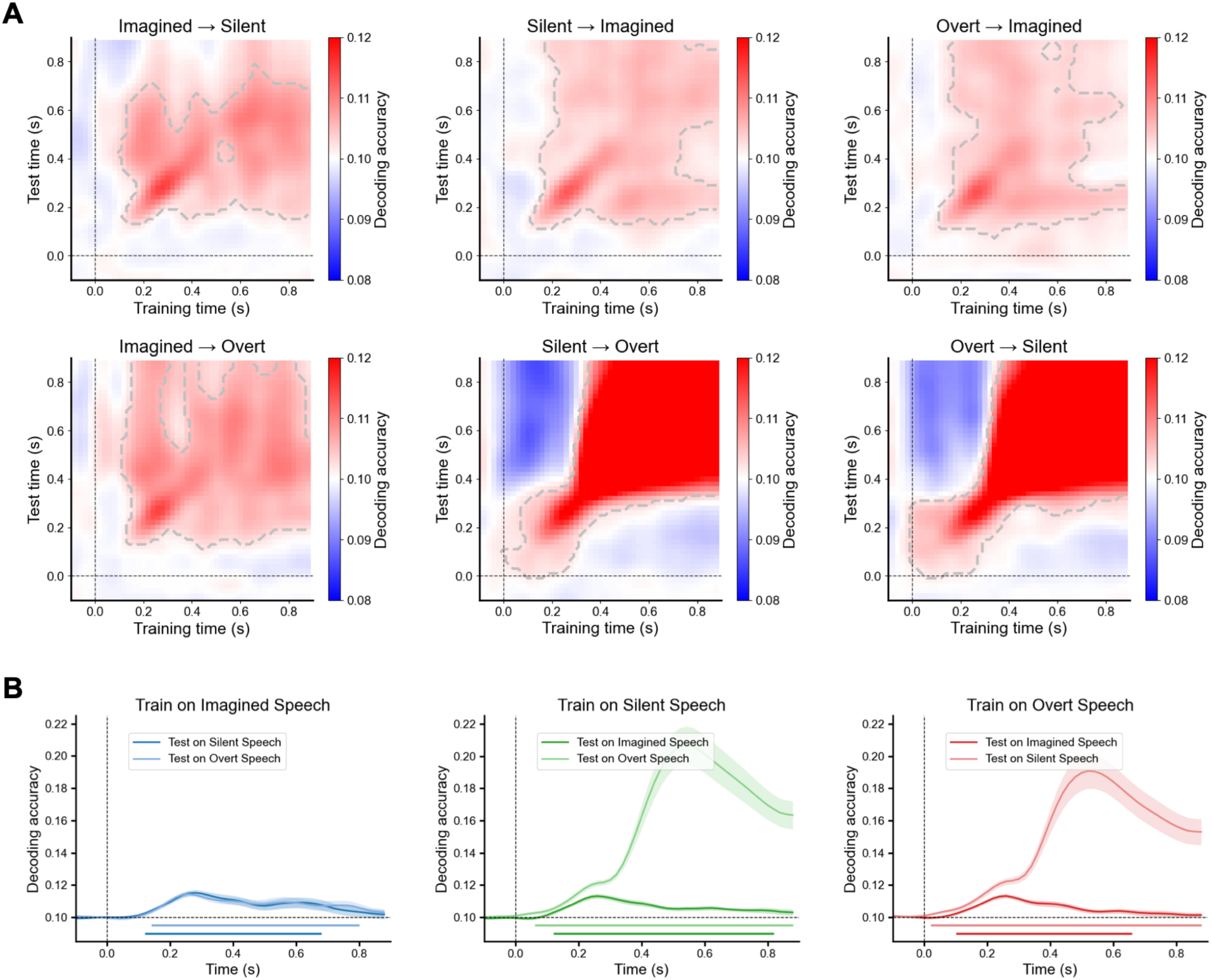
Direction-specific cross-mode decoding of Chinese word representations. (A) Unidirectional cross-mode temporal-generalization matrices for all six training-to-test transfer directions: imagined→silent, silent→imagined, overt→imagined, imagined→overt, silent→overt, and overt→silent speech. For each participant, a classifier trained to discriminate word identity at each time point in one speech mode was tested across all time points in another mode. Colors indicate the across-participant mean decoding accuracy. Horizontal and vertical black dashed lines indicate stimulus onset (0 s), and gray dashed contours delimit significant above-chance clusters identified using one-sided two-dimensional cluster-based permutation tests (chance = 0.10; 1,000 permutations; cluster-level *p* < 0.05). (B) Main-diagonal decoding time courses corresponding to the six directional transfer analyses in panel A, grouped according to the training speech mode. Curves show the across-participant mean decoding accuracy and shaded regions indicate the SEM. The horizontal dashed line indicates chance-level performance (chance = 0.10), the vertical dashed line indicates stimulus onset, and colored horizontal segments indicate significant above-chance temporal clusters identified using one-sided cluster-based permutation tests (1,000 permutations; cluster-level *p* < 0.05).

**Figure S2.**
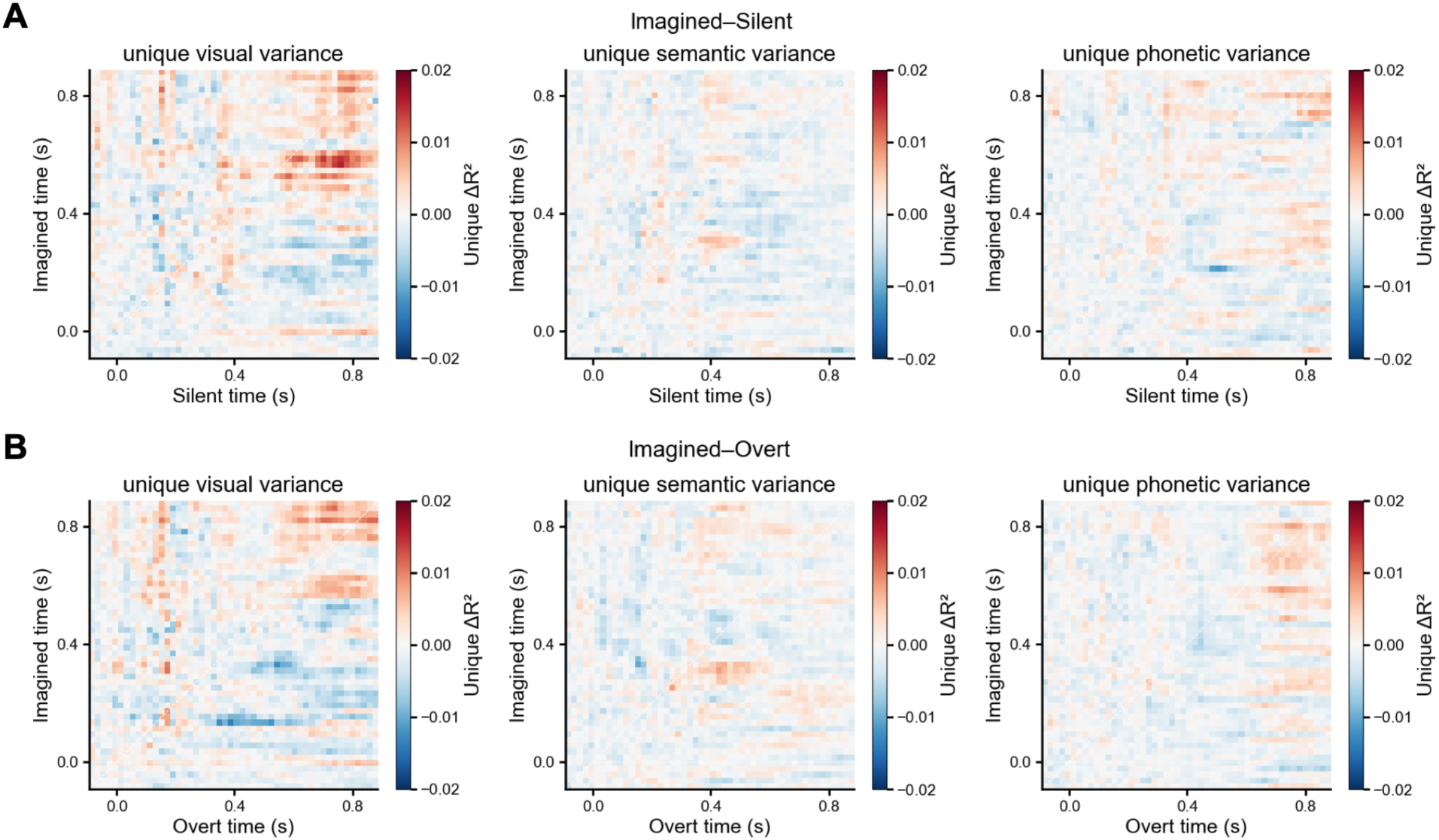
Feature-specific contributions to cross-mode representational geometry involving imagined speech. (A) Unique contributions of visual-form, semantic, and phonetic information to cross-mode EEG representational similarity between imagined and silent speech. (B) Corresponding analyses for imagined and overt speech. For each target feature, EEG RDM vectors from both speech modes were first residualized with respect to the other two feature models. Unique ΔR² was calculated as the difference between the squared residual cross-mode EEG-RDM correlation before and after additionally residualizing both EEG RDMs with respect to the target feature. Positive ΔR² values indicate that removing the target feature reduces cross-mode similarity, whereas negative values indicate an increase in residual similarity after removal of that feature. Statistical significance was assessed using two-sided two-dimensional cluster-based permutation tests against zero (1,000 permutations; cluster-level *p* < 0.05; correction performed separately for each map). No significant clusters were observed for either imagined– silent or imagined–overt speech.

